# Hippocampal Signal Complexity and Rate-of-Change Predict Navigational Performance: Evidence from a Two-Week VR Training Program

**DOI:** 10.1101/2024.03.27.587026

**Authors:** Jason D. Ozubko, Madelyn Campbell, Abigail Verhayden, Brooke Demetri, Molly Brady, Yadurshana Sivashankar, Iva Brunec

**Author notes:** Corresponding Author: Jason D. Ozubko.

## Abstract

The hippocampus is believed to be an important region for spatial navigation, helping to represent the environment and plan routes. Evidence from rodents has suggested that the hippocampus processes information in a graded manner along its long-axis, with anterior regions encoding coarse information and posterior regions encoding fine-grained information. Brunec et al. (2018) demonstrated similar patterns in humans in a navigation paradigm, showing that the anterior-posterior gradient in representational granularity and the rate of signal change exist in the human hippocampus. However, the stability of these signals and their relationship to navigational performance remain unclear. In this study, we conducted a two-week training program where participants learned to navigate through a novel city environment. We investigated inter-voxel similarity (IVS) and temporal auto-correlation hippocampal signals, measures of representational granularity and signal change, respectively. Specifically, we investigated how these signals were influenced by navigational ability (i.e., stronger vs. weaker spatial learners), training session, and navigational dynamics. Our results revealed that stronger learners exhibited a clear anterior-posterior distinction in IVS in the right hippocampus, while weaker learners showed less pronounced distinctions. Additionally, lower general IVS levels in the hippocampus were linked to better early learning. Successful navigation was characterized by faster signal change, particularly in the anterior hippocampus, whereas failed navigation lacked the anterior-posterior distinction in signal change. These findings suggest that signal complexity and signal change in the hippocampus are important factors for successful navigation, with IVS representing information organization and auto-correlation reflecting moment-to-moment updating. These findings support the idea that efficient organization of scales of representation in an environment may be necessary for efficient navigation itself. Understanding the dynamics of these neural signals provides insights into the mechanisms underlying navigational learning in humans.

## Introduction

Successful spatial navigation relies on the integration of two complementary forms of spatial representation, a concept that finds support in the structural and functional differentiation along the hippocampus’s long axis. Researchers have argued that the anterior regions of the hippocampus are responsible for encoding coarser, more generalized spatial information, while the posterior regions process more fine-grained information (Brunec et al., 2018; Patai et al., 2019; Poppenk, Evensmoen, Moscovitch, & Nadel, 2013; Sekeres, Winocur, & Moscovitch, 2018; Strange, Witter, Lein, & Moser, 2014). This granularity of information processing along the hippocampal axis is believed to support spatial navigation by ensuring that individuals can not only identify their overarching spatial objectives but also navigate the intricacies of their immediate environment to achieve these goals.

While initial evidence for this model of spatial processing has predominantly derived from rodent studies, recent research by Brunec et al. (2018) has presented evidence in humans that both signal granularity and the rate of signal change in the hippocampus vary along this axis. In their study, participants navigated through a familiar real-world city using Google Street View-like software while in an fMRI scanner. The researchers were then able to calculate the inter-voxel similarity (IVS) across voxels in both the anterior and posterior hippocampus, providing a measure of representational granularity or complexity. Similarly, the researchers calculated temporal auto-correlation during navigation in the anterior and posterior regions to obtain a measure of rate of signal change. Brunec and colleagues (2018) discovered that, similar to prior rodent findings, humans showed coarse-to-fine grained spatiotemporal representations along the long-axis of the hippocampus, from the anterior to the posterior regions respectively.

Building on Brunec and colleagues (2018) work, we present a further exploration of the dynamics of both signal complexity and signal change, as they relate to navigational performance and specifically how these signals change with experience. Studies have shown that the hippocampus exhibits dynamic changes in neural activity over the course of long-term learning. For example, during initial learning, the hippocampus may undergo rapid changes in synaptic plasticity, which gradually stabilize over time as memories consolidate (Frankland & Bontempi, 2005; Wiltgen, Brown, Talton, Silva, & Neurobiology, 2004). Brunec et al. observed that although IVS was related to navigational strategy, it was consistently higher in the anterior than the posterior hippocampus, even during periods of rest. This begs the natural question as to how stable IVS is, and whether it can change as one becomes more experienced in navigating an environment or whether its dynamics can predict navigation performance (i.e., the ability to navigate novel routes in a learned environment). As for temporal auto-correlation, Brunec and colleagues (2018) found an anterior-posterior distinction only during navigation, and not during rest. We are interested in further understanding when and how this anterior-posterior distinction in auto-correlation would emerge during navigation, whether it would change over time with gradual learning, and whether it could predict successful navigational performance.

To explore the dynamics of IVS and auto-correlation, we conducted a two-week training study in which participants were trained to navigate 4 inter-connected routes in a city that was new to them. Over the two week period, participants engaged in 5 training sessions of guided navigation and frequent quizzes in an effort to teach them the routes. From this, a group of Strong and Weak Learners emerged. Strong Learners took part in an unguided navigation day after training, where they freely navigated through several navigational challenges. (Note that pilot testing had revealed that not every participant could learn the routes in the two-week period we provided. We thus filtered participants during the experiment, allowing only those with strong performance by Session 5 to take part in Session 6, activation navigation. Participants who were able to take part in Session 6 were thus classified as Strong Learners. Those that were filtered out or that tried Session 6 but had have their sessions terminated before any useful data was collected were classified as Weak Learners). Afterwards, all participants took part in one last day of guided navigation along the 4 trained routes. Over the course of this study we scanned participants on their second day of training, and during the unguided navigation day and the final day of guided navigation. We thus gathered imaging data from early training (guided) and late training (unguided and guided). Using this data we present an exploration of both IVS and auto-correlation, and the dynamics between the anterior-posterior hippocampus. Our goal is to explore how IVS or auto-correlation signals are influenced by navigational ability (Strong vs. Weak Learners), by training (Early vs. Late sessions), and by the dynamics of navigation themselves (i.e., kind of navigation or success vs failure).

## Materials and Methods

### Participants

Thirty-six healthy right-handed volunteers were recruited. Five participants were excluded due to difficulties with the task and an inability to complete the navigational training. Of the remaining thirty-one participants who completed the study there were 6 males and 28 females (mean age 20.17 years, range 18-24 years). Of the thirty-one participants, 20 would be later identified as Strong Learners and 11 as Weak Learners (see Experimental Design).

All participants were free of psychiatric and neurological conditions. All participants had normal or corrected-to-normal vision and otherwise met the criteria for participation in fMRI studies. Informed consent was obtained from all participants in accordance with SUNY Geneseo and the University of Rochester’s ethical guidelines. Participants received monetary compensation upon completion of the study.

### Surveys and Demographics

Participants completed a brief survey to gather demographic data and assess their navigational abilities before participating in the study. This survey included (1) a question asking participants about how many hours per week they play video games; (2) the Navigational Strategies Questionnaire (Brunec et al., 2018); and (3) the Santa Barbara Sense of Direction Scale (Hegarty, Richardson, Montello, Lovelace, & Subbiah, 2002).

### Routes and Navigation Software

Researchers selected the area of Squirrel Hill, PA for this study (see Figure 1A) because it contained a fairly grid-like layout of streets. This gave participants a reasonable opportunity to develop spatial maps where they might be able to discover or infer shortcuts between routes, and because it contained a number of visually distinct sub-neighbourhoods, ranging from houses, shops, large fields and golf courses, and even streets with distinctly colored cobblestone. The variation in environments was desired as it was hoped this would help participants orient themselves as they learned the layout of all of these distinct regions.

**Figure 1.**
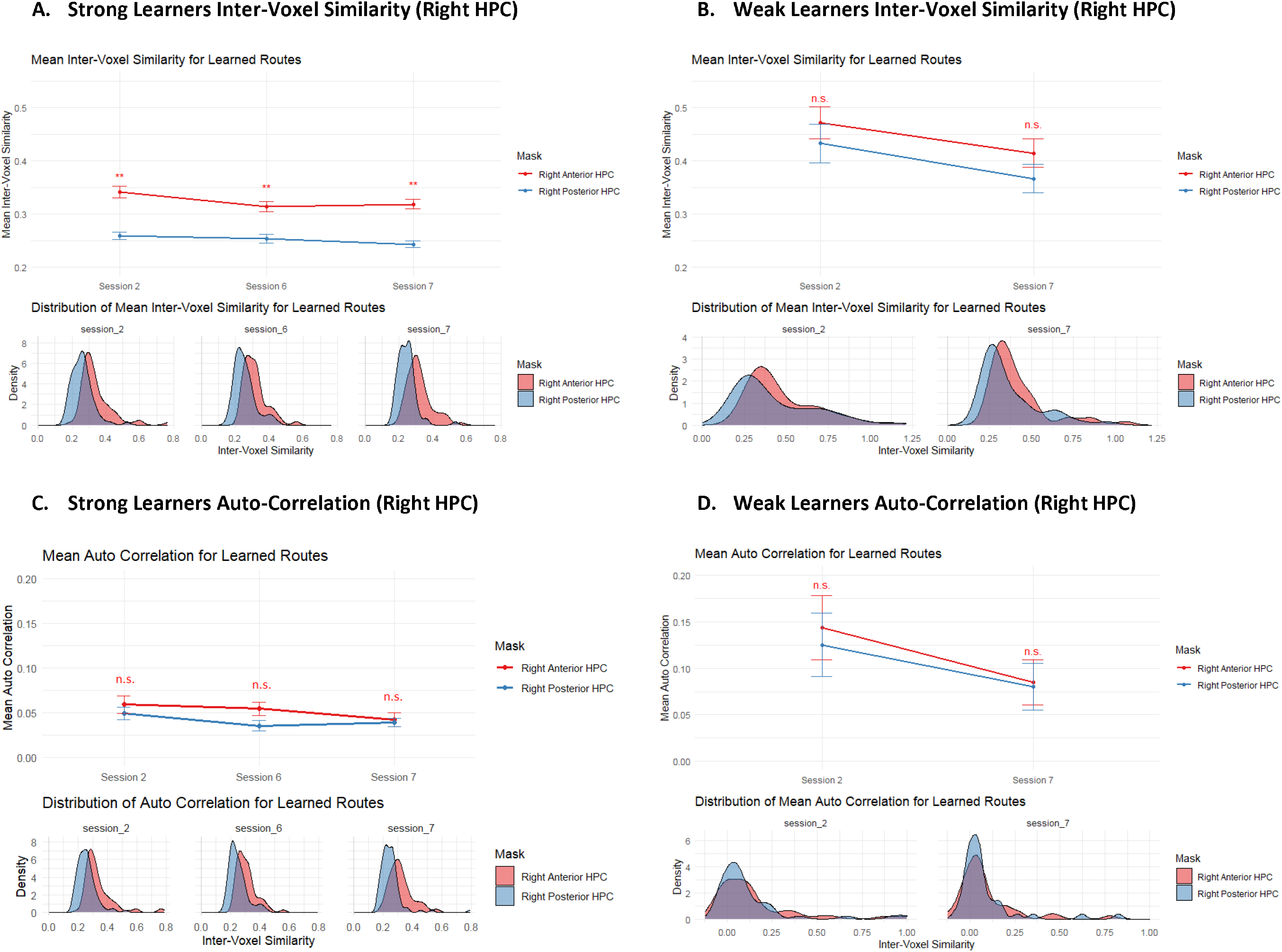
Mean inter-voxel similarity ratings and temporal auto-correlation derrived from right hippocampal activity. Results are separated by session and by learned type (Strong vs. Weak Learners). Strong Learners showed an anterior-posterior difference in inter-voxel similarity across all sessions, and in auto-correlation in Session 6. Weak Learners did not show any anterior-posterior difference in either measure and had generally highler levels of inter-voxel similarity and auto-correlation than Strong Learners. Note that Session 2 and 7 were passive navigation (following an arrow along routes) whereas Session 6 was active navigation (navigating the learned routes from memory). Only Strong Learners participated in Session 6 necessarily as Strong Learners were defined as those participants who could navigate from memory after training. Error bars represent standard error of the means. Distributions plotted below line charts represent distribution of raw scores.

A custom software suite was designed in Python to both download images from Google Street View and recreate the Google Street View experience in the lab, to allow participants to virtually navigate and walk through environments. With this software, we could place participants at any position in the neighbourhood and provide them with a panoramic, first person-view of their position. Participants could use the arrow keys (on non-scanner day) or button box buttons (on scan days) to rotate left or right in 360 degrees. They also had a button to step forward so that they could traverse the environment.

This software was used to download the Squirrel Hill, PA neighbourhood ahead of time and to run the navigation portions of the experiment.

### Experimental Design and Statistical Design

The experiment took place over seven sessions in a 2-week period (see Figure 1B). Sessions 2, 6, and 7 were scanning sessions, whereas the remaining sessions occurred outside the scanner. During the first five sessions, participants were trained to navigate 4 routes (Figure 1A) across 5 training sessions of guided navigation (with the second session being a scanning session, the rest were held outside the scanner; Figure 1B). The four learned routes were relatively non-overlapping but all passed through a central intersection in the middle of town. Each route in the training sessions began and ended at a distinctive landmark and these landmarks were used to generate names for each route. The landmark names were invented by the researchers to be distinctive and informative. The route names were: Grey’s Manor to O’Connor Country Club, Colfax Academy to Cobblestone Way, Red Brick Road to Homewood Estate, CVS Market to the Uptown Lot. At the beginning of each training route these names were presented to participants, along with an image of their destination landmark. During navigation, the name of the destination landmark was present at the top of the screen along with an arrow which directed participants where to go (see Figure 1C). At periodic intersections, the experiment would pause and participants were asked which way they would go next in order to proceed on this route. Participants could answer left, right, or forward, and were immediately presented with feedback via the arrow updating in the correct direction after their response. Accuracy for these directional quizzes is presented in Figure 1F. Following each training trial a set of recognition tests were presented where participants were shown scenes from the route they navigated and had to identify whether the scenes came from the route at hand or were being presented in the correct order.

For each participant, the four training routes were randomly assigned to be Routes 1, 2, 3, and 4. In the first session of training, only Routes 1 and 2 were practiced (see Figure 1B). Session 2 was the first scan and all 4 routes were trained during this session. Hence, Routes 1 and 2 were recently experienced whereas Routes 3 and 4 were completely novel, providing us with data on both kinds of routes in this early scan. In Sessions 3, 4, and 5, Routes 1-4 were repeatedly practiced, prior to navigating in the scanner (in Sessions 6 and 7). Each training session took approximately 1.5-2 hours and 11 routes were practiced during each session.

Participants who exhibited steady learning in these conditions were classified into the Strong Learners category and were invited to participate in the sixth experimental session (an unguided/free navigation session). Strong Learners showed 80% or greater accuracy on directional quizzes by Session 5 (see Figure 1F). Participants who exhibited difficulty with the navigational task and who showed poorer learning performance were classified as Weak Learners and not invited to take part in the sixth experimental session. In pilot testing participants who had not learned the routes well floundered during the sixth experimental session, which involved various navigational challenges, and often became frustrated. Given these issues and the fact that we also had budgetary constraints, we allocated the sixth day scans to the Strong Learners only.

The specific trials in the sixth session included Route Following, Route Finding, and Blocked Routes. For Route Following, participants were asked to travel each of the four training routes accurately from memory. For Route Finding, participants were placed at one of the starting landmarks and asked to navigate to the destination landmark from a different route, thus requiring them to find a way to bridge between two known routes. Lastly, for Blocked Routes participants were asked to travel a trained route from memory, however, they were informed that the center of town had been blocked off and that when they reached the obstruction they would have to find an alternate route to the destination. We used the experimental software to place artificial road signs around the center square of town and thus physically prevented participants from traversing those roads, forcing them to find alternate routes to their destination. In total, there were 4 Route Following trials, up to 9 Route Finding trials, and 4 Blocked Route trials. Participants had a maximum of 8 minutes to navigate any particular route but in practice were not given 8 minutes on every route due to scanner timing limitations—if participants took too long navigating on some trials (or had clearly become lost) the researcher had to skip other trials in order to keep the experiment on track. Thus, due to these complex timing balances, we prioritized collecting all 4 Route Following and all 4 Blocked Route trials, and then also collecting as many Route Finding trials as time would allow.

Finally, in Session 7 participants were presented with each of the four trained routes once, and were guided along the route in the exact same manner as they were in the first five training sessions. This final session was carried out in the scanner so that we could compare guided navigation in Session 2 (early in training) with guided navigation in Session 7 (after training). Statistical design for this Experiment can be found in the following fMRI Image Acquisition section as well as the Results section.

### fMRI Image Acquisition

Scanning took place at the University of Rochester Center for Advanced Brain Imaging & Neurophysiology. Participants were scanned on a Siemens 3T Prisma MRI scanner. On the first scan day (the second experimental session) a high-resolution 3D MPRAGE T1-weighted pulse sequence image (192 axial slices, 1 mm thick, FOV = 256 mm) was first obtained to register functional maps against brain anatomy. Functional imaging was performed to measure brain activation by means of the blood oxygenation level dependent (BOLD) effect. Functional T2*-weighted images were acquired using echo-planar imagine (84 axial slices, 2 mm thick, TR = 1500 ms, TE = 30 ms, flip angle = 60 degrees, FOV = 200 mm). The native EPI resolution was 128 x 128 with a voxel size of 2mm x 2mm x 2mm.

Images were preprocessed using the FEAT tool in FSL v6.00. Preprocessing involved motion correction (MCFLIRT), brain extraction (BET), spatial smoothing (5mm kernel), high-pass filtering, spatial realignment, and co-registration. Pre-processed data was then transformed into MNI space.

Following preprocessing, we employed methods to quantify the similarity of activation patterns across different brain regions and their stability over time, specifically focusing on IVS and temporal autocorrelation, as delineated by Brunec et al. (2018). To calculate IVS, we first computed the Pearson correlation coefficients between the BOLD signal time series of all pairs of voxels within predefined anterior and posterior ROIs in the hippocampus. This process was facilitated by constructing voxel-by-voxel correlation matrices for each participant, reflecting the degree of functional connectivity between voxels during navigation tasks. The mean correlation coefficient across all voxel pairs within an ROI was taken as the measure of IVS, indicative of the coherence of neural representations within that spatial region.

Temporal autocorrelation was assessed by calculating the correlation of the BOLD signal time series of each voxel with its own lagged series across the entire navigation task. This was achieved by shifting the time series by one TR (1500 ms) and computing the Pearson correlation for each voxel with its lagged self. This measure reflects the stability or persistence of the neural signal over time, providing insight into the temporal properties of neural encoding during spatial navigation. Both IVS and temporal autocorrelation metrics were computed for each participant and then averaged across the group to obtain mean values. All analyses were conducted using custom scripts in R and Python, alongside the FSL toolbox.

## Results

To preface and summarize our results, we find that Strong Learners exhibited a clear anterior-posterior distinction in IVS and some anterior-posterior distinction in auto-correlation in the right hippocampus, while Weak Learners showed less pronounced distinctions. However, lower overall IVS levels in both the anterior and posterior right hippocampus were linked to better early learning. As for auto-correlation, successful navigation was characterized by faster signal change, particularly in the anterior hippocampus, whereas failed navigation lacked the anterior-posterior distinction in signal change. These findings suggest that signal complexity and signal change in the hippocampus are important factors for successful navigation, with IVS representing information organization and auto-correlation reflecting moment-to-moment updating.

### Behavioural Measures: Strong vs. Weak Learners

These behavioural data have been reported previously in Ozubko et al. (under review). We will summarize only the major points here. Participants who exhibited steady learning the training phase (Sessions 1 through 5) were classified into the Strong Learners category and were invited to participate in the sixth experimental session (an unguided/free navigation session). Strong Learners showed 80% or greater accuracy on directional quizzes by the end of Session 5. Participants who struggled with the navigation task and demonstrated poorer learning performance were identified as Weak Learners and were consequently excluded from the sixth experimental session. From pilot testing, we observed that those who were less proficient in learning the routes struggled significantly in this session. The sixth session introduced a variety of navigational challenges, which often led to frustration among the Weak Learners. Due to these challenges, coupled with budget constraints, we decided to reserve the scans on the sixth day for those identified as Strong Learners only.

Strong and Weak Learners did not differ on weekly hours spent playing video games, *t*(29) = 0.51, *p* =.61, navigation strategies, *t*(29) = 0.78, *p* =.44, sense of direction, *t*(29) = 0.36, *p* =.72, or age, *t*(29) = 1.00, *p* =.33. However, Strong Learners significantly outperformed Weak Learners on navigational directional quizzes, achieving an average accuracy of.90 (SE =.02) after Session 5, in contrast to the Weak Learners’ average accuracy of.70, *t*(29) = 4.48, *p* <.01. It’s worth mentioning that Weak Learners were relatively adept at navigating directions, but they were not as precise as Strong Learners. Indeed, Strong Learners consistently showed higher accuracy than Weak Learners in directional quiz accuracy across all training sessions, except the first, all *p*’s <.05. Interestingly, Strong Learners were slower to respond to directional quizzes than were Weak Learners during the first training session, *t*(29) = 2.37, *p* <.05, although this difference quickly subsided with further training. By the final session, Strong Learners displayed a tendency to be quicker than Weak Learners on directional quizzes, though not significantly so, *t*(29) = 1.81, *p* =.09. However, Strong Learners were significantly quicker in navigation during the last training session, *t*(29) = 2.49, *p* <.05. No other significant differences in navigation times were observed in any other training sessions. In summary, Strong Learners distinguished themselves through more accurate navigational learning, with only slight advantages in navigation speed and no differences in overall navigation skills or abilities.

### Signal Complexity, but not Rate-of-Change, is related to Navigational Performance

As mentioned previously, IVS is used as a measure of signal complexity and auto-correlation as a measure of rate-of-change. Mean IVS and auto-correlation values for anterior and posterior regions of the right hippocampus, separated by session and learner type (Strong vs. Weak) and session (Sessions 2, 6, and 7) can be seen in Figures 1. Our goal in these analyses is to assess the influence of experience (i.e., differences between sessions) and skill (i.e., differences between Strong and Weak Learners) on IVS and auto-correlation.

Strong Learners showed an anterior-posterior distinction in IVS across all sessions, with higher levels of IVS in anterior than posterior regions (Figure 1A; all *t*’s > 3.49, *p*’s <.01). Consistent with prior research, this result suggests that Strong Learners are representing more complex (finger-grained) representations in posterior regions than anterior. Interestingly, this anterior-posterior IVS distinction is not only absent in Weak Learners (Figure 1B; both *t*’s < 1, *p*’s >.42), but Weak Learners exhibit higher overall levels of IVS than Strong Learners across all sessions (both *t*’s > 2.34, *p*’s <.05). It can be inferred then that Weak Learners may not be processing information complexity in as distinct a manner along the long-axis of their hippocampus and may generally be representing less complex representations than Strong Learners. Interestingly, though non-significant, IVS levels numerically decline for Weak Learners from Session 2 to Session 7 suggesting that these learners *may* gradually be adapting through experience to represent slightly more complex representations (both *t*’s < 1, *p*’s >.39), albeit still not as complex as Strong Learners.

Auto-correlation levels of Strong Learners can be seen in Figure 1C and of Weak Learners in Figure 1D. There was no anterior-posterior difference in auto-correlation levels observed in any session for either Strong, all *t*’s < 1.03, *p*’s >.31, or Weak Learners, both *t*’s < 1, *p*’s >.84. As with IVS, Weak Learners once again demonstrated higher levels of auto-correlation than Strong Learners however this difference was also non-significant, both *t*’s < 1.13, *p*’s >.28. In short, the rate of signal change, as measured by auto-correlation, did not different significantly along the hippocampus between Strong and Weak Learners or across sessions.

Briefly examining the left hippocampus for reference, the same general patterns observed for the right hippocampus emerged, albeit the anterior-posterior distinction was more muted for Strong Learners (See Appendix A). For Strong Learners, the anterior-posterior difference in IVS was only marginal across Sessions 2, 6, and 7, all *t*’s > 1.81, *p*’s <.08. All other patterns from the right hippocampus replicated, suggesting that while the complexity of representations along the long-axis may not have been as clearly defined in the left hippocampus, Strong Learners still showed more complex representations compared to Weak Learners, and may have still been differentiating complexity along the long-axis better than Weak Learners. For auto-correlation there were no significant values to report in the left hippocampus.

Overall, these data demonstrate that an anterior-posterior distinction in the right hippocampus may contribute to superior spatial learning and navigation, particularly in regards to representational complexity. High overall levels of IVS coincide with poor performance, suggesting that overall levels of IVS and auto-correlation must remain relatively low for optimal performance, and differentiated along the long-axis of the hippocampus. In our remaining analyses we will examine in more detail how IVS in the right hippocampus is tied to navigational performance.

### Complex Signals predict Rapid Acquisition of Novel Environments

In Session 2, we measured both across-session and within-session learning. In Session 1 of the experiment, subjects were guided along Route 1 six times and Route 2 twice. In Session 2, participants were once again guided along these two routes, providing us with measures for activity during route guidance for that had been recently exposed across session. Two new routes were also shown in Session 2—Routes 3 and 4— and repeated several times. Participants were guided during navigation on Route 3 four times and Route 4 twice. Routes 3 and 4 thus provide measures for routes that have been learned within session. Analyses here focused on participant performance on the intersection quizzes and how it relates to IVS. Specifically, this relation lets us examine how well participants were learning the four routes, and whether their memorization of the routes related to IVS.

In Figure 2A we have the intersection quiz accuracy plotted for each trial in Session 2, with the trial sequence being: Route 1, Route 2, Route 3, Route 4, Route 3, Route 3, Route 3, and Route 4. Similarly, we are plotting the mean IVS from each of those 8 trials. Linear regression analysis was used here to contrast runs and check for differences among multiple conditions simultaneously. As can be seen in Figure 2A, quiz performance was best for Route 1 compared to all other routes, *B* =-0.27, *SE* = 0.04, *t*(60) = 28.48, *p* <.01. Since Route 1 had been exposed 6 times in Session 1 (more than any other route), this result was expected. Route 2 was also exposed in Session 1 (though only twice), and thus showed lower quiz accuracy than Route 1, *B* =-0.22, *SE* = 0.22, *t*(60) = 5.33, *p* <.01, but better quiz accuracy than the first presentation of Routes 3 and 4 (i.e., trials 3 and 4), *B* =-0.17, *SE* = 0.03, *t*(60) = 6.18, *p* <.01. Route 3 and Route 4 were shown for the first time on trials 3 and 4 respectively and showed lower quiz accuracy on their first exposure than on their final exposure (trials 7 and 8), *B* =-0.20, *SE* = 0.03, *t*(60) = 5.95, *p* <.01.

**Figure 2.**
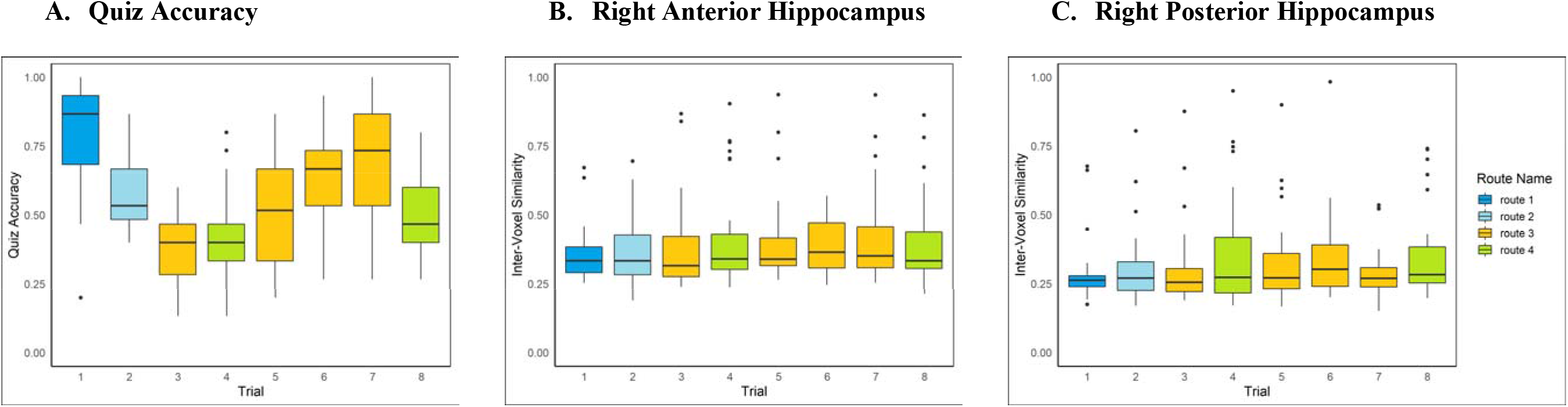
Quiz accuracy, right anterior hippocampus inter-voxel similarity, and right posterior hippocampus inter-voxel similarity for the 8 trials from Session 2. Each boxplot shows the median (central line), interquartile range (IQR, edges of the box), and 1.5x IQR (whiskers) to illustrate the distribution spread. Outliers beyond the whiskers are indicated with dots.

In Figure 2B we have IVS levels for each of the 8 trials plotted individually. One-way ANOVAs were carried out as an omnibus check for significance between trials here. While quiz performance did vary substantially across the trials, mean levels of IVS did not significantly differ across any of the trials in either the anterior or posterior right hippocampus (both *F*’s < 0.47, *p*’s >.85). These data thus tell a relatively simple story, which is that knowledge of routes increased in proportion to exposure but overall IVS levels were unaffected by exposure. Although overall IVS levels may have been stable, the more interesting question is whether there any relation between quiz performance and IVS levels.

Figure 3 plots the correlation between IVS and quiz performance for each of the 8 trials. Regression models were fit separately to anterior and posterior IVS levels and mean quiz accuracy for each subject. Results of these analyses are reported in the upper right corner of each scatterplot. As can be seen in the plots, IVS did not correlated with quiz performance except on trials 5 and 8, all R2’s >.184, p’s <.05. These represent the first repetition of Routes 3 and 4 respectively. That this relation did not persist for Route 3 on its subsequent repetitions (i.e., trials 6 and 7) indicate that during very early learning, lower levels of IVS in both the anterior and posterior hippocampus were associated with better performance on the intersection quizzes.

**Figure 3.**
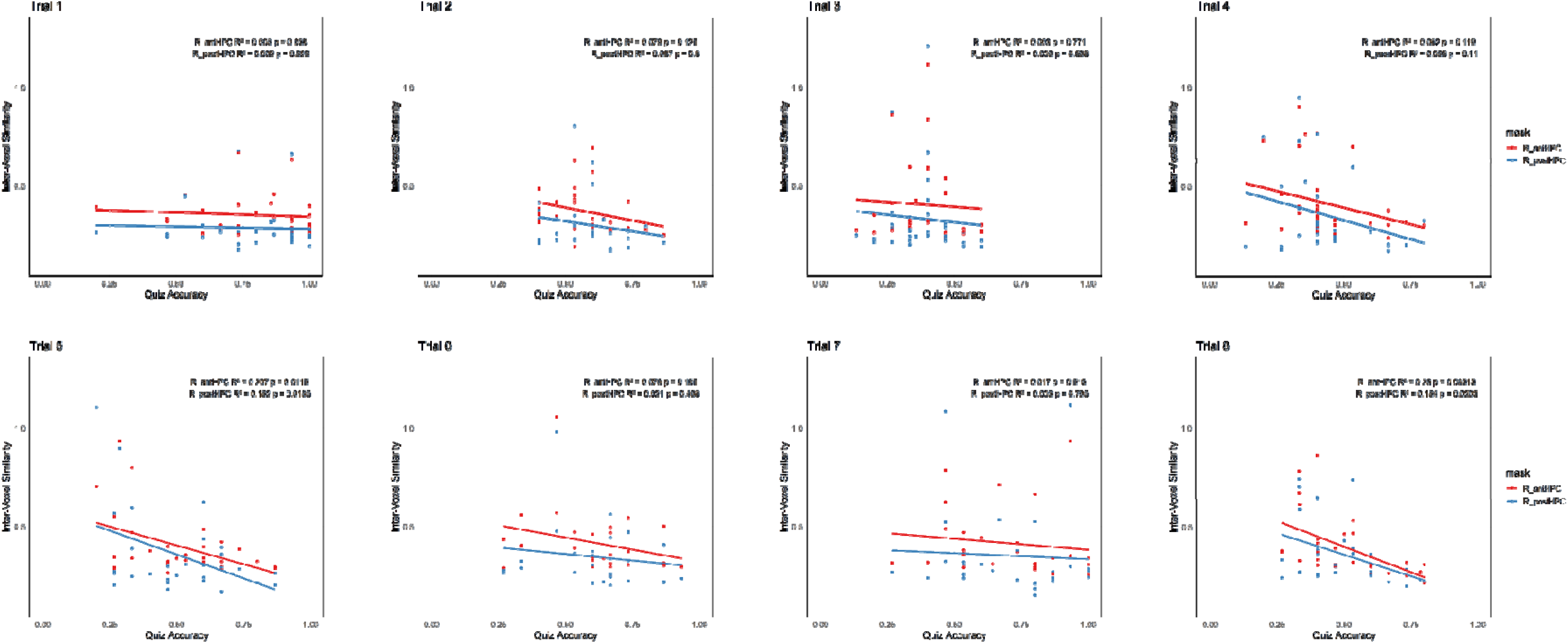
Relations between quiz accuracy and right anterior or right posterior hippocampus inter-voxel similarity ratings for the 8 trials from Session 2. Each scatterplot represents a different trial from Session 2. Each point represents data from a single subject, with red points showing the inter-voxel similarity values from the right anterior hippocampus and blue points showing the inter-voxel similarity values from the right posterior hippocampus. Lines shown are the regression lines between inter-voxel similarity and quiz accuracy across subjects.

That lower IVS levels helped early learning, we believe, is indicative of how complexity in representations is valuable early when learning a route. Indeed, low levels of IVS may represent accurate mapping of space into early mental maps. Why then would IVS not continue to correlate with route performance? As subjects continue to learn routes, there are likely numerous strategies that could yield accurate performance on the intersection quiz, ranging from spatial strategies like cognitive maps to rote visual strategies such as memorizing some particular landmark in the background for each given turn. Given that a variety of strategies could help and subjects were likely using an array of strategies in a task as complex as ours, low levels of IVS may have helped subjects who were using a mapping strategy gain an early advantage, but as everyone gradually caught up, the necessary importance of IVS faded and thus the correlations became nonsignificant in later repetitions (i.e., trials 6 and 7), all *R^2^’s <.076, p*’s *>*.17.

Overall, these data highlight that an important relation does exist between complexity of representation and acquisition of novel environments, but that this relation exists briefly, in the earliest stages of learning.

### Signal Complexity Predicts Failures after Blockages, Signal Stability Predicts Route-Retracing Success

Thus far, signal complexity (as measured by IVS) has differentiated Strong from Weak Learners and been linked to early route learning, indicating an overall connection between IVS and route learning, but a specific connection between individual route performance and IVS has not been clear. Here we explore the possibility that whereas signal complexity provides a gross measure of navigational learning, signal stability can provide a more fine-grained picture of performance on individual routes. Specifically, we explore how temporal auto-correlation in the right hippocampus differentiates successful from unsuccessfully navigated routes in Session 6. Once again by focusing solely on Session 6 we are exploring only the data from Strong Learners. If anything though, this should make it more difficult to find significant differences as the distinction between Strong Learners is smaller than the distinctions between Strong and Weak Learners.

Figure 4 plots the IVS and auto-correlation for Repetition Trials, Cross-Overs, and Blocked Trials in Session 6. Critically, we are plotting successful trials (trials where subjects reached the goals; marked a “1”) and failed trials separate (failed data has the dimmer colors; marked as “0”). For IVS, the anterior-posterior difference is present in all trial types almost regardless of performance, all *t*’s > 3.70, *p*’s <.01, once again contributing to the notion that IVS provides a gross measure of navigational processes but may not be sensitive to route-specific happenings. The one exception is the blocked routes, *t*(19) = 0.92, *p* =.37, which we will explore further below. For auto-correlation, the anterior-posterior difference may be more closely tied to overall performance during route-retracing as on Repetition Trials a marginal effect is present on successful trials, *t*(19) = 2.01, *p* =.05, but in all other cases no anterior-posterior difference was observed, all *t*’s < 1.74, *p*’s >.09. These data suggest that it is stable representations that, on an individual trial basis, may be required for successful route-retracing.

**Figure 4.**
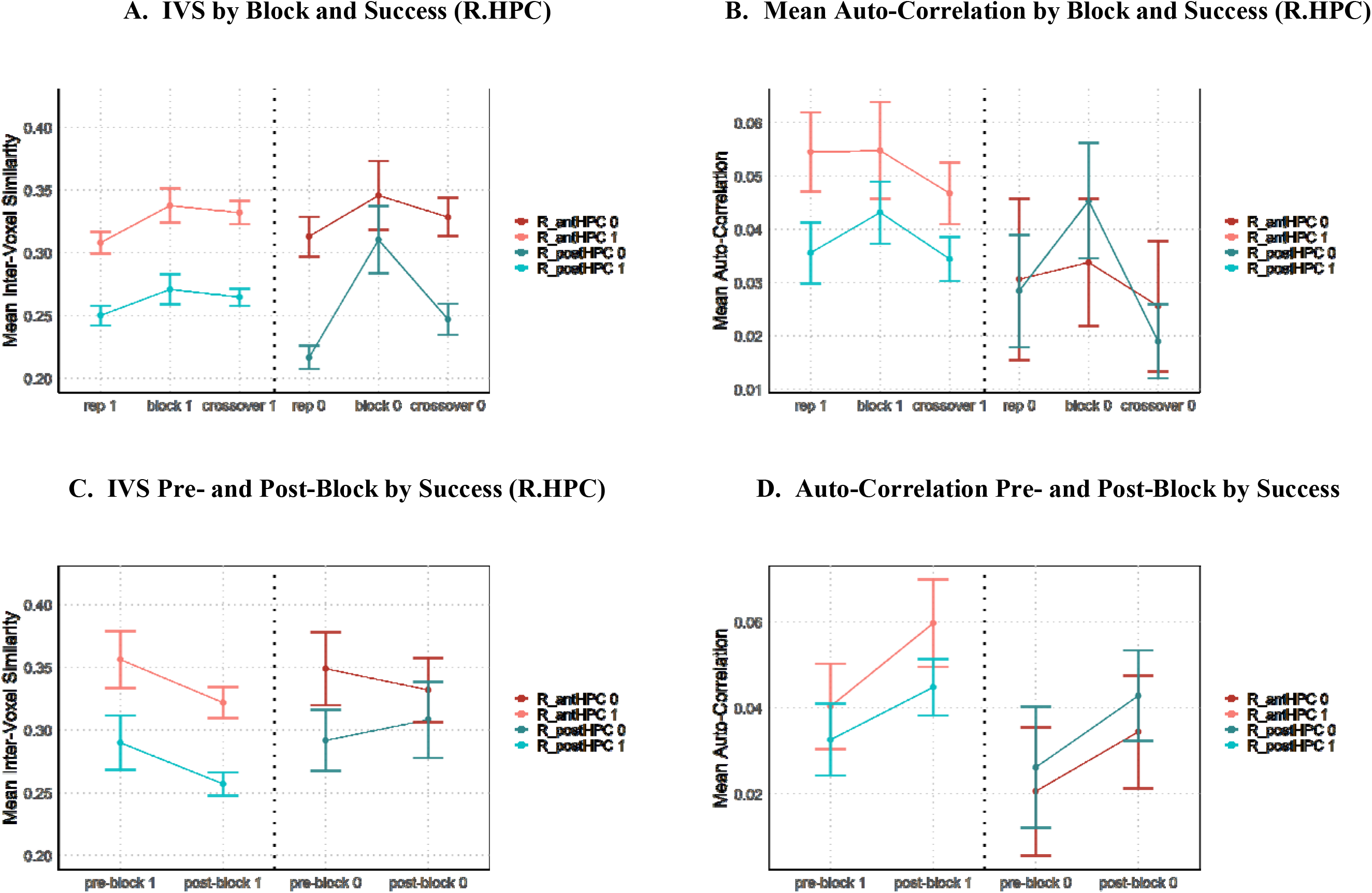
Mean IVS and auto-correlation levels for right anterior and posterior hippocampus. Panels A and B show mean levels from the right hippocampus for Repetition Trials, Cross-Over Trials, and Blocked Trials, separating success (bright, left-side) from failed (dim, right-side). Panels C and D show mean levels from the right hippocampus from Blocked Trials before and after the blockage was encountered, separating success (bright, left-side) from failed (dim, right-side).

Exploring Block Trials just a bit more, there is one more notable pattern to report. On successfully navigated block trials, participants show an anterior-posterior distinction in IVS both prior to encountering the blockage and afterwards, both *t*’s > 2.30, *p*’s <.05. However, on failed trials (i.e., trials where participants ultimately got lost and could not find the goal), the anterior-posterior distinction in IVS is present pre-blockage, *t*(12) = 2.68, *p* <.05, but not post, *t*(12) = 2.04, *p* =.06. (Note that the reduced degrees of freedom for these tests are a result of not every subject having failed at least one blocked route). Furthermore there were no significant anterior-posterior differences between auto-correlation values either pre-or post-block as a function of trial success, all *t*’s < 0.28, *p*’s >.79. As the anterior-posterior IVS distinction has been tied to the usage of maps already, we interpret this finding as showing that navigational failures occur when individuals cannot maintain their mental map post-blockage. In other words, these data are showing attempts to navigate around the blockage that renders one lost and unable to place oneself on ones mental map. Thus, while representational stability may generally predict navigational success, the ability to coherently maintain a mental map can be vital in some circumstances and directly predict performance on a trial-by-trial basis.

## General Discussion

We sought to explore the how signal complexity and signal stability contribute to navigational success in a long-term learning paradigm. Our results show that higher levels of signal complexity in the right hippocampus are tied to superior navigational abilities and the usage of mental maps. More so, there is a necessary anterior-posterior distinction, with superior navigational abilities demonstrated when the posterior right hippocampus showed more complex signals than the anterior. Indeed, failure to differentiate along the long-axis resulted in more difficulty learning routes and difficulty navigating around obstacles along routes. Higher levels of signal complexity also contributed to better early learning of routes. As for signal stability, faster signal change in the right hippocampus generally indicated better performance, albeit within limits. An anterior-posterior distinction was once again observed where, in optimal situations, signals should change more slowly in the anterior than the posterior right hippocampus. Indeed, the anterior-posterior distinction was not observed on trials which participants failed to navigate. In sum, whereas signal complexity and signal change represent distinct aspects of representation during navigation, both are important for successful navigation and the signals within the right hippocampus appear particularly relevant.

Our results align with previous observation that higher levels of signal complexity in the hippocampus are associated with better early learning (Brunec et al., 2018) and studies showing that the richness and complexity of neural representations can facilitate learning and memory formation (Leutgeb, 2005; Pilly & Grossberg, 2013). Other research has shown that neural representations of space can change as environments become well learned (Hirshhorn, Grady, Rosenbaum, Winocur, & Moscovitch, 2012; Maguire, Nannery, & Spiers, 2006; Patai et al., 2019; Rosenbaum et al., 2000; Spiers & Maguire, 2008; Teng & Squire, 1999; Wolbers & Wiener, 2014). Though our research was necessarily limited to a shorter time-frame than these past studies which have examined the difference between remotely-learned and recently-experienced environments, we nonetheless found evidence that even very early learning can produce unique patterns in the hippocampus compared to later learning. Namely, our observation that the first repetition of routes in Session 2 elicited a correlation between IVS in the right hippocampus and performance suggests that when environments are being learned, complex representations in the hippocampus may be crucial to facilitate accurate acquisition. These findings suggest that the ability of the hippocampus to represent fine-grained information is crucial for effective learning and accurate spatial navigation.

Strong navigators show more complex and faster changing representations than Weak learners. This is true early in training, late training, and whether it’s unguided or guided navigation. The finding that we did not observe any significant changes in pattern complexity whether the navigation was guided or unguided (i.e., free active exploration) reveals that individual differences related to navigational learning matters more than the type of navigation engaged at encoding. For example, researchers have found that the presence of a navigational aid (i.e., analogous to guided navigation) impairs spatial memory assessed by recall of landmarks, and accuracy in later map drawings (Gardony, Brunyé, Mahoney, & Taylor, 2013). However, the current findings suggest that a key element underlying navigational performance is the anterior-posterior distinction in the hippocampus. In terms of IVS, this distinction is more broadly present, consistent with the idea that the hippocampus may have inherent structural biases to process information broadly in anterior and in fine-grained manner in posterior. Nonetheless, Weak Learners did not show this pattern as clearly as Strong Learners. Failure to represent complexity in this graduated manner along the long-axis might therefore contribute to poor learning. As well, poor learners generally showed less complexity and more slowly changing signals. This improved with training but likely also could have contributed to their poorer ability to learn the environments.

For Strong Learners, during unguided navigation the anterior-posterior distinction was present in IVS regardless of performance except on Blocked Routes. On Blocked routes the anterior-posterior distinction was disrupted by the blockage on Routes that would ultimately be failures. It’s possible that on these routes participants had difficulty maintaining coherent cognitive maps as they deviated from their known route. When maps could be maintained, the anterior-posterior distinction was held intact and navigation was a success. When participants became lost, this anterior-posterior distinction and map coherence deteriorated. However, this explanation is tentative considering that we did not observe an anterior-posterior breakdown on Route Finding (i.e., Cross-Over) trials.

As for signal change, auto-correlation was clearly related to navigation performance. Increased signal change, especially in the anterior portion of the hippocampus, eliminated the anterior-posterior difference and was associated with failed navigation. This increased signal change may represent an inability to identify one’s location with respect to their cognitive maps, which leads to a continued effortful updating of one’s map, as one attempts to figure out one’s location. In contrast, when subjects were successful their cognitive maps changed more slowly. Overall then these data suggest that when freely navigating the rate of signal change in the hippocampus must find an optimal level; too much or too little updating are both sub-optimal: Whereas under-updating may be indicative that one is not keeping up with one’s location in space, over-updating may represent one’s inability to find one’s location in mental space, and a continued mental effort to do so.

In sum, our results support past arguments that IVS may represent the organization of information in the hippocampus, such as the granularity of representation and the strategies one employs. Alternatively auto-correlation represents an in-the-moment picture of how information is being updated in the regions. While IVS is related to whether you are a successful navigator generally, auto-correlation is more directly related to the performance on specific trials and the effectiveness of moment-to-moment navigation. The anterior-posterior hippocampal distinction between both of these signals appears to be important for proper route learning and navigation. This critical distinction that we observed primarily in Strong relative to Weak Learners highlights the important role of individual differences pertaining to navigational performance.

## Acknowledgments

We wish to thank Harris Schwab, Kaitlyn Bertleff, and Luke Bamburoski for help carrying out this experiment. We also wish to thank the staff at the University of Rochester Center for Advanced Brain Imaging & Neurophysiology for their support. This research was supported by NIH (1R15NS104979-01).

## Notes

### Competing Interest Statement

The authors have declared no competing interest.

