## Appendix for "Hippocampal Signal Complexity and Rate-of-Change Predict Navigational Performance: Evidence from a Two-Week VR Training Program"

**Appendix A:**

**Analyses of IVS and Auto-correlation in the Left Hippocampus**

In complementary analyses to our main analyses, we extended our examination to the left hippocampus to ascertain whether the patterns observed in the right hippocampus were consistent across hemispheres. Specifically, we focused on IVS and auto-correlation measures across anterior and posterior regions, differentiated by learner type (Strong vs. Weak) and by session (Sessions 2, 6, and 7). The aim was to investigate potential hemispheric differences in signal complexity and rate-of-change representation.

The results, presented in the accompanying figure (see Figure S1), reveal that the general patterns identified in the right hippocampus largely mirror those in the left, albeit with some distinctions. For Strong Learners, the anticipated anterior-posterior gradient in IVS is present but less pronounced, with marginal t-values suggesting a trend but not reaching conventional levels of statistical significance, all *t*’s > 1.81, *p*’s < .08. This suggests a subtle, yet not robust, differentiation in complexity representation along the long axis of the left hippocampus among Strong Learners.

Furthermore, for both Strong and Weak Learners, no significant differences were found in auto-correlation measures across the anterior and posterior regions of the left hippocampus (all t’s < 1, p’s > .28). This parallels findings in the right hippocampus, underscoring the non-significant differentiation in the rate of signal change between learner types and across sessions.

In sum, the left hippocampus showed a reduced anterior-posterior distinction in IVS across sessions for Strong Learners than the right hippocampus. All other anterior-posterior differences were nonsignificant.

| A. **Strong Learners Inter-Voxel Similarity (Left HPC)**   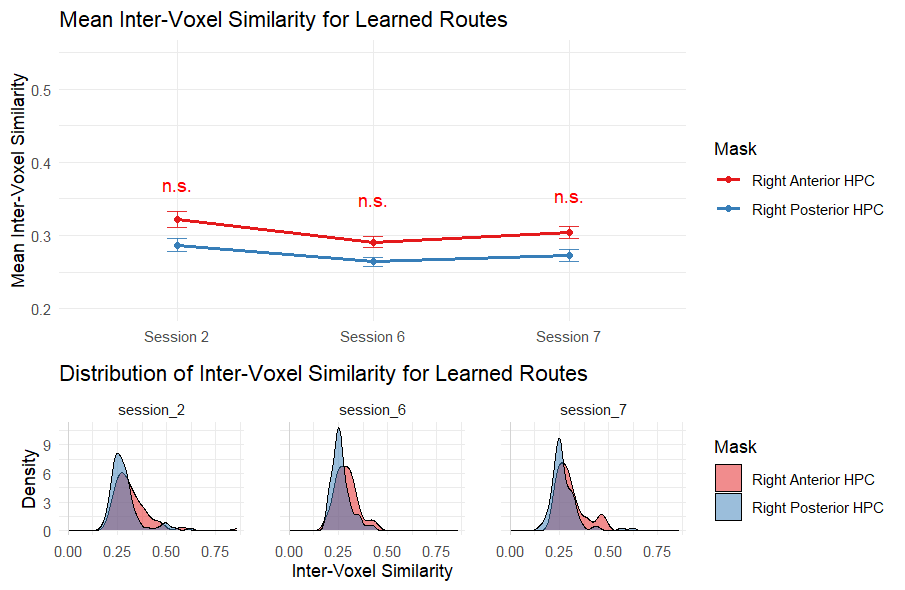 | B. **Weak Learners Inter-Voxel Similarity (Left HPC)**   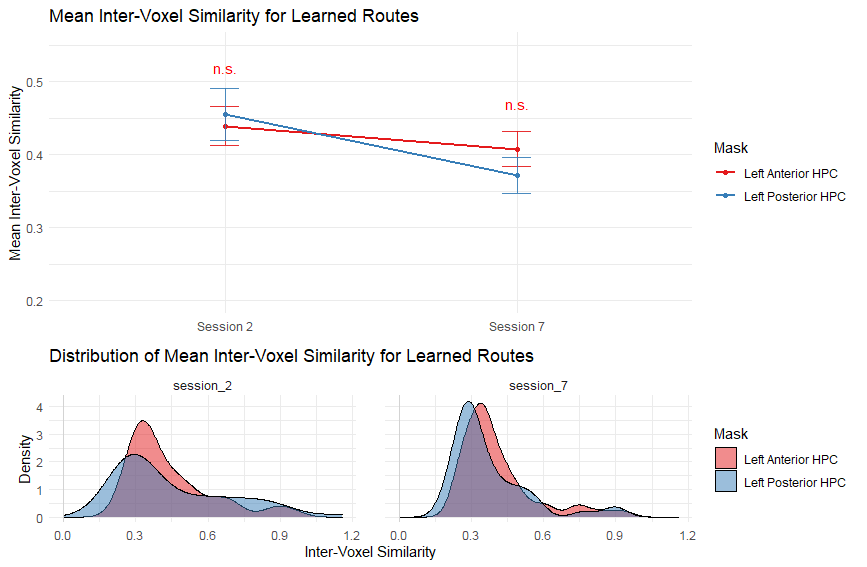 |
| --- | --- |
| C. **Strong Learners Auto-Correlation (Left HPC)**   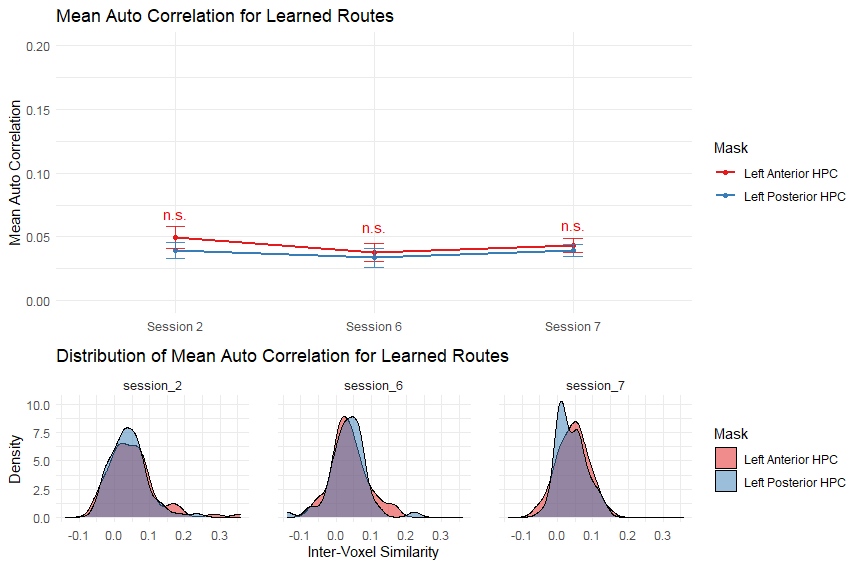 | D. **Weak Learners Auto-Correlation (Left HPC)**   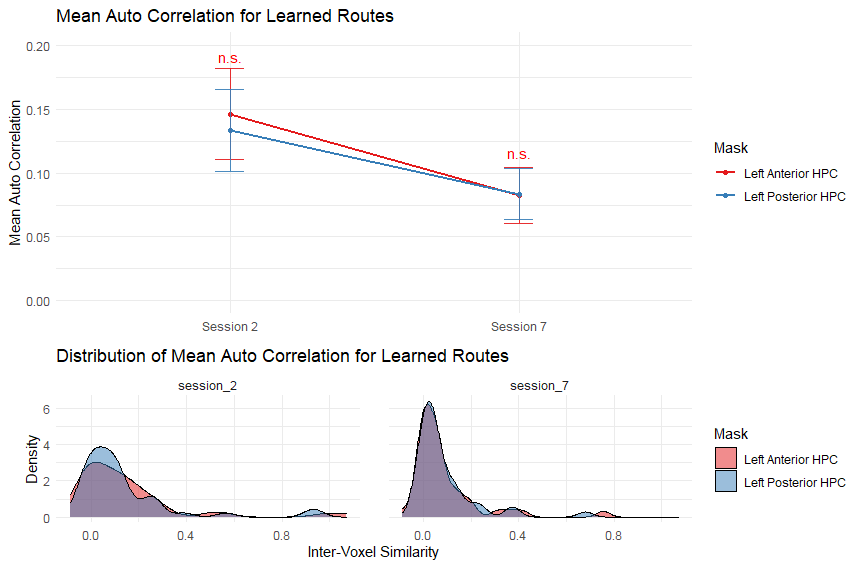 |

**Figure S1.** Mean inter-voxel similarity ratings and temporal auto-correlation derrived from left hippocampal activity. Results are separated by session and by learned type (Strong vs. Weak Learners). No anterior-posterior difference was observed for any condition for Strong or Weak Learners in either inter-voxel similarity or auto-correlation measures. Error bars represent standard error of the means. Distributions plotted below line charts represent distribution of raw scores.
